# STrack: A tool to Simply Track bacterial cells in microscopy time-lapse images

**DOI:** 10.1101/2022.12.20.521337

**Authors:** Helena Todorov, Tania Miguel Trabajo, Jan Roelof van der Meer

## Abstract

Bacterial growth can be studied at the single cell-level through time-lapse microscopy imaging. Technical advances in microscopy lead to increasing image quality, which in turn allows to visualize larger areas of growth, containing more and more cells. In this context, the use of automated computational tools becomes essential.

In this paper, we present STrack, a tool that allows to track cells in time-lapse images in a fast and efficient way. We compared it to three recently published tracking tools on images ranging over six different bacterial strains, and STrack showed to be the most consistent tracking tool, returning more than 80% of correct cell lineages on average.

The python implementation of STrack, a docker structure, and a tutorial on how to download and use the tool can be found on the following github page: https://github.com/Helena-todd/STrack

## 1 Introduction

Despite major advances in methodologies describing microbial systems’ states (e.g., the typical ‘o-mics’ tools), there is a fundamental aspect of microbial behaviour that is escaping much of our attention. This concerns *in situ* cell division, phenotypic heterogeneity, cell movement and consequently, local population growth. For most microbiome systems, individual taxa cannot be easily studied in their natural biotic context, nor followed in real-time. Most information, therefore, comes from experiments of reduced complexity, where cell growth can be followed directly by microscopy time-lapse imaging (Kron [2002]). Time-lapse imaging provides direct information on bacterial cell shapes, sizes, and division rates, as well as more complex phenotypes, such as cell movement or stabbing, at the single-cell level. It also allows to visualise the organisation of cells into colonies or spatial structures that result from food intake or from interspecific interactions. Relevant information on cell division rates, spatial processes or interactions between bacteria can be derived from the images resulting from time-lapse experiments. This requires to identify single cells in the images (resulting in what is called “segmented masks”), and to track these masks across all time steps correctly.

Several methods have been recently published to tackle the complex task of tracking cells across time-lapse images (Jeckel and Drescher [2021], Ulman et al. [2017]). Some of these tools integrate both cell segmentation and tracking in a single pipeline (Meacock et al. [2021], Versari et al. [2017], Stylianidou et al. [2016]). Since cell segmentation often leads to errors that might have a big impact on the subsequent tracking steps, the tools typically provide a graphical user interface (GUI) that allows users to manually curate results before applying cell tracking. Identified cells are then linked in successive frames based on cell-to-cell distances and similarities. The tracking algorithms are not error prone, and the GUI also offers the possibility to correct tracks manually. Highly interactive methods can thus help to obtain perfect tracking results through extensive curation, but they present the drawback of requiring a lot of investment from the users.

Automated tracking tools that can take segmented masks as input have also been recently introduced. For example, Trackmate (Tinevez et al. [2017]) has been used to track particles (Fazeli et al. 2020, Omelchenko et al. [2021]), and can take object shapes into account when assigning tracks (Ershov et al. [2021]). A newer tracking method, Trac^X^ (Cuny et al. [2021]), additionally considers the environment of a cell to decide which cells should be linked in successive frames. A third method, DeLTA 2.0 (O’Connor et al. 2022), relies on two neural networks to identify and track cells in an automated way. These three methods are automized, but they can require extensive fine-tuning from the user (in the case of Trac^X^), specific Python scripting (e.g., DeLTA), or be embedded in commercial softwares such as Matlab (e.g., Trac^X^ and SuperSegger).

The goal of the work presented here was to simplify automated bacterial cell tracking from time-lapse imaging. The tool we developed and tested (called *STrack*, for *S*implified cell *Track*ing) is implemented in the free Python programming language and runs using one single line of code. We intentionally reduced the parameters that the user needs to define to a minimum of two intuitive parameters. The first one defines a restricted search space around a cell, by which the tracking algorithm finds corresponding cells in subsequent images, which drastically reduces the computational running time of STrack. The second parameter defines the division axis, which allows to improve tracking of rodshaped bacteria specifically, by allowing them to divide only along their cell elongation axis. To facilitate the use of STrack on any operating system, we wrapped it in a docker structure (Boettiger [2015]). The Docker system prevents conflicts with previously installed packages on one’s computer, and makes STrack’s results more reproducible, as it will return exactly the same results regardless of any updates of the libraries it relies on.

In the first section of this paper, we introduce the STrack algorithm in detail. Then, in section 2, we present the time-lapse datasets on which we applied STrack, as well as the methods that we used to assess the accuracy of different cell tracking tools. Section 3 is a comparison of STrack to three of the most recent automated tracking tools: TrackMate, Track^X^ and DeLTA 2.0 (see above), on the same image sets. To compare the tracking performance of these tools, we used expert-generated manual tracking results as ground-truth. We found that STrack consistently outperformed the other three automated methods on an average over the 22 datasets that we used. Furthermore, we also assessed the tools’ accessibility from a user’s point of view by comparing practical features such as their running time, free availability, and number of parameters. Finally, in the last section of this paper, we discuss STrack’s advantages and limitations.

## 2 Methods

### 2.1 Algorithm

We designed STrack to track segmented objects, that result from manual or automated single-cell segmentation, between successive timepoint images. We adapted the tool to optimise tracking of rod-shaped and coccoid bacterial cells growing in planar conditions with single cell layers, such as produced in microfluidic devices (Dal Co et al. [2020]) or surface growth chambers (Reinhard et al. [2010]). Bacterial cells typically divide by elongation, with an elongated mother cell (Timepoint 0, Figure 1, A and C) giving birth to two smaller daughter cells (Timepoint 1, Figure 1, B and D). STrack requires only two parameters, which relate directly to the images, to be set by the user:

**Fig. 1:**
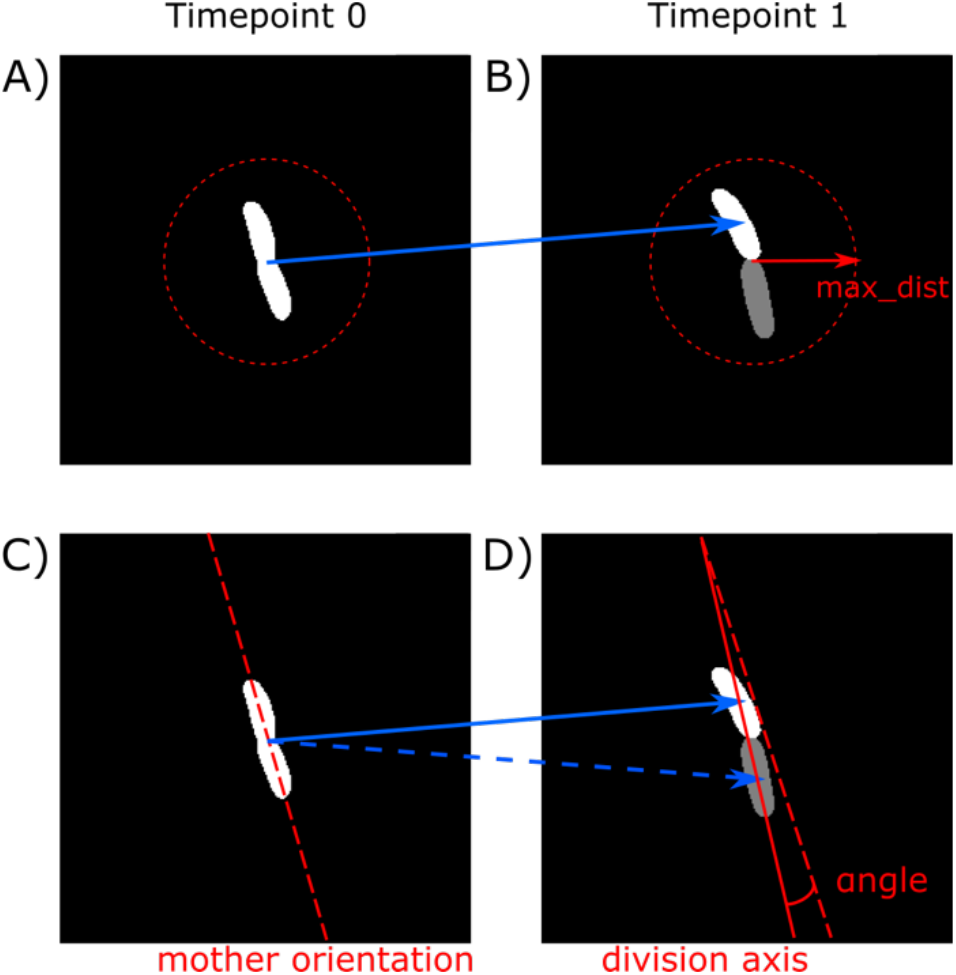
STrack requires two user-defined parameters. The first parameter max_dist restricts the search space around a mother cell (A) to look for daughter cells (B). The second parameter max_angle limits the angle between the mother cell’s main orientation axis (C), and the division axis passing through the centres of its two putative daughter cells (D). If the angle between these two axes is smaller than the user-defined threshold, then STrack will assign tracks (represented as blue arrows) between the mother cell and its two daughter cells.

- the maximum distance (max_dist) used to look for the progeny of a mother cell (see Figure 1, A and B). Setting a maximum distance drastically reduces the time and memory STrack takes to run, rendering it applicable to images containing hundreds of cells.
- the maximum angle (max_angle) formed by a mother cell and its two daughter cells (Figure 1, C and D). This prevents STrack from allowing a mother cell to divide along an axis that would be perpendicular to its elongation axis, which is less plausible from a biological point of view.

The algorithm’s pseudocode is provided in Algorithm 1. It iterates over all images of a time-lapse. For every set of two successive timepoints, the algorithm will compute distances and pixel-overlap between objects in the first image (we call them mother cells) and objects in the second image (daughter cells) that are in a perimeter delimited by the *max_dist* parameter. Based on the pixel overlap, the algorithm will then assign mother-daughter links (starting with the maximum pixel overlap and then decreasingly looking for cells with less overlap). For a second daughter cell to be assigned to a mother cell, the angle between the mother cell’s main axis and the division axis needs to be smaller than the user-provided *max_angle* parameter.

Once all mother-daughter links have been assigned based on pixel overlap, there may remain cells with no mother cells. At this point, the algorithm will switch from pixel overlap to distances between cells to identify any potential missing tracks. It will assign mother-daughter links for increasingly larger mother-daughter cell distances (while still respecting the constraints on the maximum division angle defined above). If the distances become larger than the user defined *max_dist* parameter, the remaining cells will be assigned to new tracks. This allows to start tracking cells that enter the image in the middle of a time-lapse.

The output of STrack is a single CSV-formatted table for every timepoint. These tables contain tracks from mother cells to daughter cells, including their respective X and Y coordinates, thus allowing to manually verify whether tracks were correctly assigned. STrack also returns one image per timepoint, in which the links from the previous to the current image are shown. This facilitates visual checking of the results, as one can directly see whether tracks were correctly assigned between cells or not (Fig.2).

**Fig. 2:**
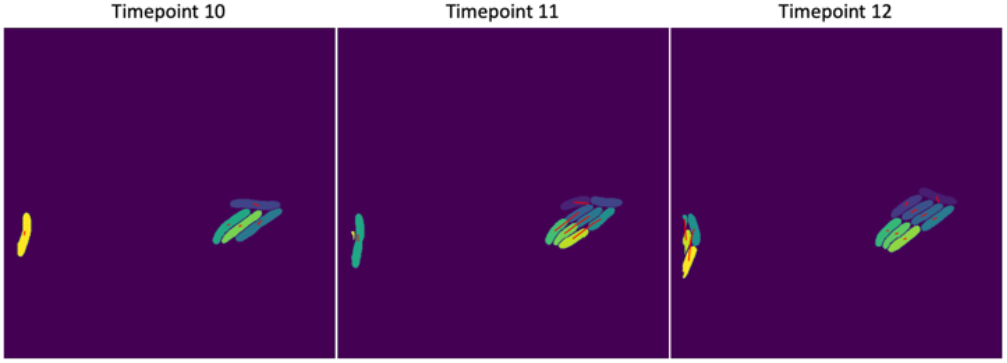
Visual cell tracking validation. For each timpoint, STrack outputs the original cell-mask-image on which the cell tracks that it identified are overlayed. These tracks (represented as red lines) start in the centre of mother cells in the image corresponding to the previous timepoint, and end in the centre of their daughter cells in the current image. In this example, STrack identified five cells at timepoint 10, the majority of which divided between timepoints 10 and 11. These divisions can be seen in the image corresponding to timepoint 11 as red lines that originate from the centres of their mother cells in timepoint 10 and lead to their respective daughter cells in timepoint 11. In this example, STrack even managed to accurately identify the progeny of the two cells that were very close to the left image border in timepoint 11.

### 2.2 Datasets

#### Time-lapse imaging

Bacterial cells were grown on miniaturised nutrient agarose surfaces that allow single cells to grow into monolayer microcolonies. Such miniaturised surfaces are enclosed in a black anodised POC chamber (H. Saur Laborbedarf, Germany), which is then mounted on a Nikon ECLIPSE Ti Series inverted microscope coupled with a Hamamatsu C11440 22CU camera and a Nikon CFI Plan Apo Lambda 100X Oil objective, at 22 °C.

Time-lapse imaging was controlled by a script in MicroManager Studio (v1.4.23). Phase contrast images were taken every 10-20 minutes, depending on the strains, with an exposure time of 30 ms. Cells were imaged on 8-10 randomly selected positions per surface, for a duration of 12-20 h, resulting in .tif files. We focused on four bacterial species isolated from soil: *Pseudomonas putida, Pseudomonas veronii, Lysobacter sp*. and *Rahnella sp* (the two latter species coming from Causevic et al. [2022]). Cultures were recovered from –80 °C stocks and grown individually on nutrient agar, from which a single colony was transferred and grown in liquid media, before being washed, diluted, and inoculated on the microscale-surfaces for imaging. Both pseudomonads were grown with 1 mM succinate as sole carbon substrate in type 21C minimal medium added to the agarose (Gerhardt et al. [1981]). The other two strains were cultured with tenfold diluted PTYG-medium (peptone, tryptone, yeast extract and glucose), as described by Bakken and Olsen [1987].

#### Manual image processing

All time-lapses presented in this paper (including the two datasets derived from the literature) were analysed manually to generate ground-truth segmentation and tracking results. Experts in the field of microbiology manually defined cell masks in these images using the QuPath open-source software for bioimage analysis (Bankhead et al. [2017]). This resulted in more than a thousand segmented cells over 22 time-lapse data series. The resulting segmented cells were then manually tracked in successive images by the same experts, using the MaMut tracking Fiji plugin (Wolff et al. [2018]). This produced the set of manually extracted tracks that we used as groundtruth to compare to the results of automated tracking tools.

#### Automated image processing

In order to assess the computation time necessary for completion of the tracking task by the different tools presented in this paper, we re-used five positions of the *P*.*putida* and *Rahnella* strains, but taking into account more timepoints than the ones that had been manually annotated. We purposely selected time-lapses with high cell numbers, in crowded images (Supplementary Table 1). These datasets were used to assess the running times only, not the tracking quality itself. We segmented the cells in these images using the automated cell segmentation tool Omnipose (Cutler et al. [2022). The resulting segmented masks were given as input to STrack, Trax^X^, TrackMate and DeLTA 2.0, and their computation time was measured on a mac with a i7-9750H processor, 32Go RAM DDR4 running Catalina operating system version 10.15.7.

### 2.3 Comparison of tracking tools

#### Automated tracking tools

The cell masks resulting from manual cell segmentation (as described above) were used as input for STrack and the three other automated tracking tools that we compared it to. To track cell objects using TrackMate, we used a procedure from [2021] described by J. W. Pylvanainen on the imageJ plugins website (https://imagej.net/plugins/trackmate/trackmate-label-image-detector). We did not set any filters on the spots or the tracks, and used the following parameter values: frame-to-frame linking was set to 100 pixels, no gap closing was allowed, track segment splitting was allowed with a corresponding maximum distance of 50 pixels. In the case of Trac^X^, we contacted the authors who generously shared a script with us (see Supplementary Trac^X^ MATLAB script). For DeLTA 2.0, the authors shared a script on gitlab that we used to inject our manual segmentation results, and use only the tracking feature of DeLTA 2.0 (https://gitlab.com/dunloplab/delta/-/issues/44).

#### Comparing tracking tools

the results of STrack, TrackMate, DeLTA 2.0 and Trac^X^ were compared to the manually generated groundtruth cell lineages by quantifying matching and discordant tracks, using the Jaccard index. This index reflects the proportion of lineages that were correctly identified by a tool, among the total number of tracks present in the tool’s results and the ground-truth tracks. A Jaccard index of one would correspond to a perfect match between the tracks returned by a tool and the ground truth.

## 3 Results

To compare the performance of STrack to existing tracking tools, we grew four bacterial species in pure culture on agarose patches and imaged them using time-lapse microscopy. We limited nutrients in those patches to have a short exponential growth phase, to prevent cells from growing in multiple layers. In the resulting phase-contrast images, cells were then manually segmented by experts (Fig.3).

**Fig. 3:**
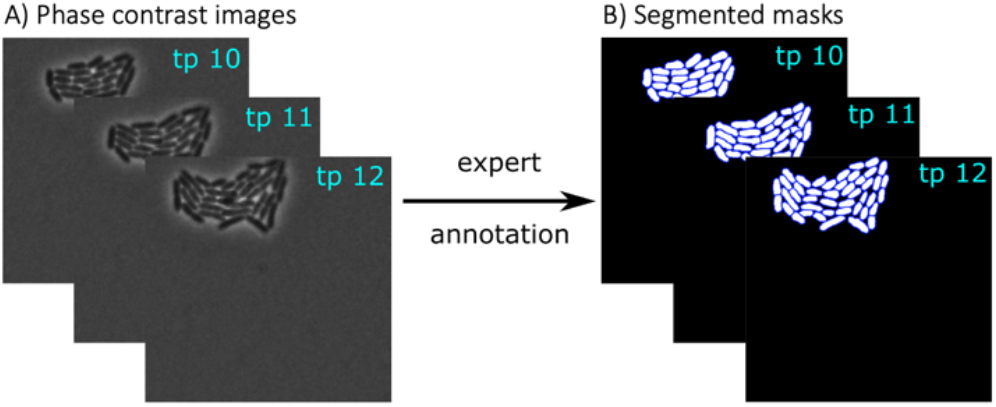
Manual expert segmentation as ground truth for STrack. A) Data consisted of phase contrast images capturing growth of bacteria into microcolonies. This figure shows an example of three timepoints (timepoints 10 to 12, each ten minutes apart) of a Pseudomonas putida microcolony. B) Identified manually segmented cells (light on dark background).

The different species displayed differences in their growth and cell shapes, with round-shaped densely packed microcolonies (i.e., *P. putida* and *Rahnella*, Fig. 4A, C), or elongated thinner microcolonies (i.e., *P*.*veronii* and *Lysobacter*, Fig. 4B and D). Two supplementary datasets from the literature were added to this study, with further differences in growth patterns that we believed could represent a challenge for cell tracking. The first dataset consists of dividing *Streptococcus pneumoniae* (Gallay et al. [2021]), a coccoid bacterium that grows into long chains (Fig. 4E).

**Fig. 4:**
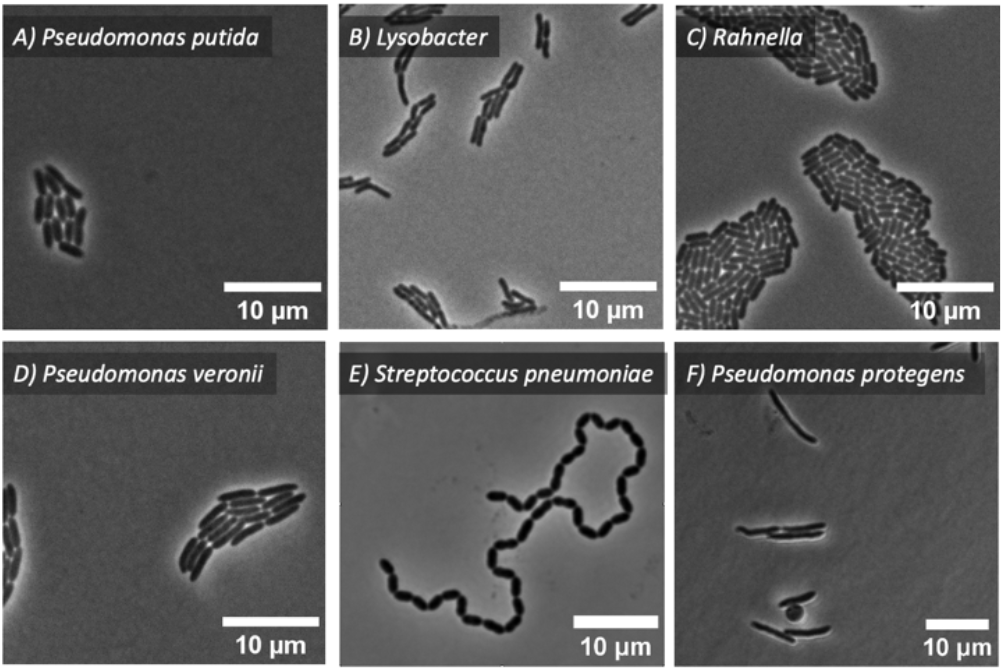
Representative images of the six bacterial strains that were included in this study, to compare how accurately STrack, TrackMate, Trac^X^ and DeLTA 2.0 would track these cells in successive frames. Note the different individual cell morphologies as well as resulting microcolony shapes. The round object in F) is a spheroblast formed by the activation of tailocins in P. protegens (Vacheron et al., [2021]). Image in E): Gallay et al, [2021].

The second showcases *Pseudomonas protegens* cell division (Vacheron et al. [2021]). The induction of R-tailocin formation in *P*.*protegens* cells results in cell elongation, followed by the formation of a spheroblast shortly before cell death (Fig. 4F).

Manually segmented cells were tracked by experts across successive images (Fig. 5A), to build the reference tracks and cell lineage trees for automated tool comparison (see Fig. 5B). The automated tracking tools were then compared using the Jaccard index, which represents the proportion of matching and discordant tracks between the results of automated tracking tools and reference tracks (Fig. 5C and 5D).

**Fig. 5.**
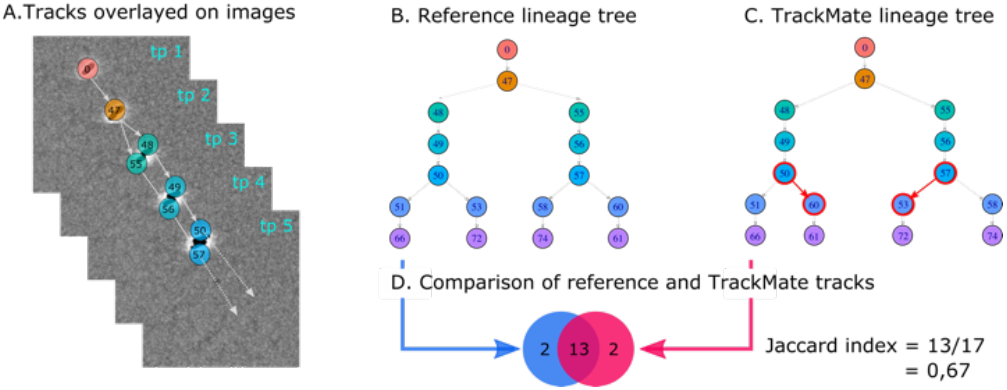
Comparison of tracking results using the Jaccard index. A) Manually tracked P. putida cells across time points (tp).The tracks are represented as grey arrows and the cells were colored and numbered to be easily compared to B) the lineage tree corresponding to the tracks from panel A. C) The results of an automated resolved tracking lineage tree over the same images, by TrackMate. Red circles and arrows highlight discordant lineages in comparison to the reference tree. D) Example of Jaccard index computation. TrackMate returned 13 similar and 2 discordant tracks, resulting in an index equal to the sum of common tracks (=13) divided by the total number of tracks in both methods (=17).

The results of the comparison of STrack and the three other tracking tools on the expert-annotated time-lapse series are shown in Figure 6. STrack and Trac^X^ significantly outperformed TrackMate and DeLTA 2.0 on the *P. putida* and *P*.*veronii* series (Fig. 6B and D), but DeLTA 2.0 had a higher average Jaccard index on the *Rahnella* images (Fig. 6C), where STrack was the second-best performing tool. Finally, STrack outperformed the three other tools in the images with *Lysobacter* (Fig. 6A). We then proceeded to a similar comparison of STrack and the three other tracking tools on two imaging datasets from the literature, consisting of a set with *P*.*protegens* images and another one with *S. pneumoniae (Fig 6E*.). All tools returned excellent automated tracking results on these two datasets, which can be explained by the fact that the *P*.*protegens* cells almost did not move or divide, and by the fact that the *S. pneumoniae* cells were imaged at very close time intervals, which greatly facilitated their tracking. STrack, DeLTA 2.0 and TrackMate had an average Jaccard index between 0.98 and 0.99 on the two literature datasets, while the Jaccard indices of Trac^X^ were slightly lower (JI = 0.931 and 0.947).

**Fig. 6:**
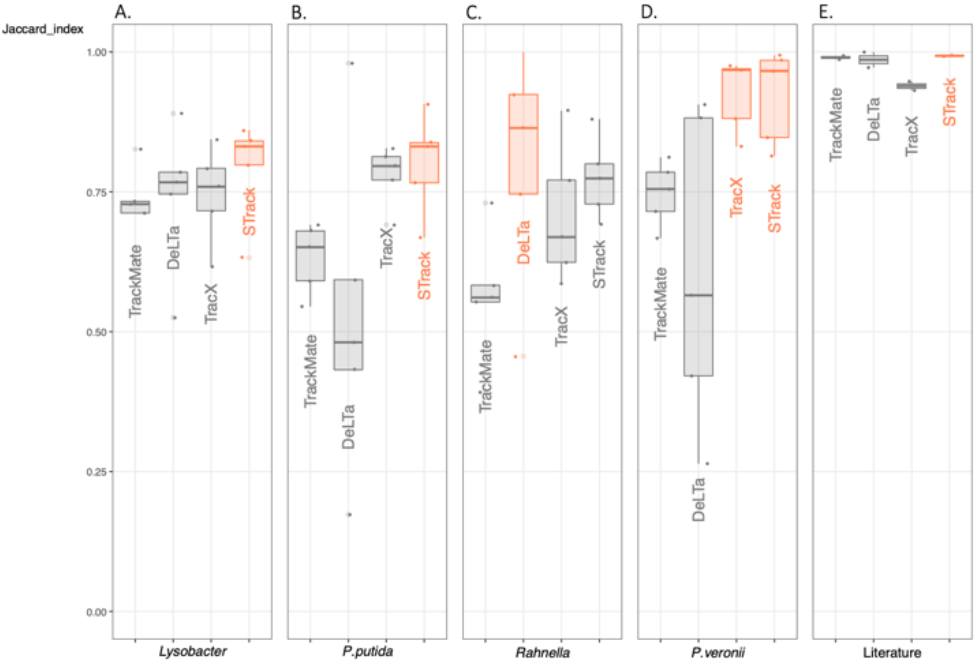
Comparison of four cell tracking tools, TrackMate, DeLTA 2.0, Trac^X^ and STrack, on microcolony growth of six different bacterial strains. A-D. Four strains were cultured and imaged in house (Lysobacter, P. putida, Rahnella and P. veronii), E. and two datasets were taken from the literature (corresponding to the strains P. protegens and S. pneumoniae). One Jaccard index was computed for each tool and for each of the twentytwo time-lapse positions. Boxplots show median (vertical line) and outlier Jaccard indices, overlaid with the individual data points. For each strain, the best performing tools (i.e., with the highest average Jaccard index) are highlighted in orange.

Across all datasets, the median STrack-JI was higher than that of the three others. STrack returned over 83% of correct tracks on average (JI = 0.838), while the second-best tool in this comparative study, Trac^X^, had an average Jaccard index of 0.803, and DeLTA 2.0 and TrackMate, had very similar scores of 0.699 and 0.696, respectively. We also noticed that the variance in individual JI’s of STrack was small, especially compared to TrackMate and DeLTA 2.0, which returned less than 50% of correct tracks on at least one dataset, while STrack consistently returned over 63% of correct tracks.

As an example of STrack’s stability compared to the other tracking tools, we show tracking results on two imaged areas of the same time-lapse of *P. putida* (Fig. 7 and 8). Although these two imaged areas come from the same time-lapse experiment, the performance of DeLTA 2.0 drastically changed between the two areas. In the first one (Fig. 7A), DeLTA 2.0 returned only 43% of matched tracks, with mismatches spanning all over the lineage tree. However, on the second dataset (Fig. 8A), DeLTA 2.0 almost perfectly matched all ground truth tracks. One obvious difference between the two datasets is that one microcolony grew very close to the image border, with some cells exiting the frame (Fig. 7B), while the other microcolony grew at the very centre of the images (Fig. 8B). We investigated possible links between the distance of microcolonies to image borders and tracking results, and found a positive correlation for DeLTA 2.0, whose accuracy clearly increased when microcolonies grew further away from borders (see Supplementary Figure 1). TracX and TrackMate were quite consistent between the two imaged areas, returning tracking mismatches that spanned over the whole lineage tree, irrespective of the microcolony position relative to the image borders. STrack also returned relatively consistent results on these two datasets, although it performed slightly worse on the dataset presented in Figure 8 (JI = 0.77) compared to the dataset in Figure 7 (JI = 0.84). In both cases, the tracking errors of STrack were mostly localised at later timepoints down the lineage trees, when cells typically become more difficult to track as the images become crowded. TrackMate systematically performed poorer compared to the other tools, perhaps because it was optimized to track objects with complex shapes (Ershov et al. [2021]), whereas the objects here are all very similar in shape and size.

**Fig. 7:**
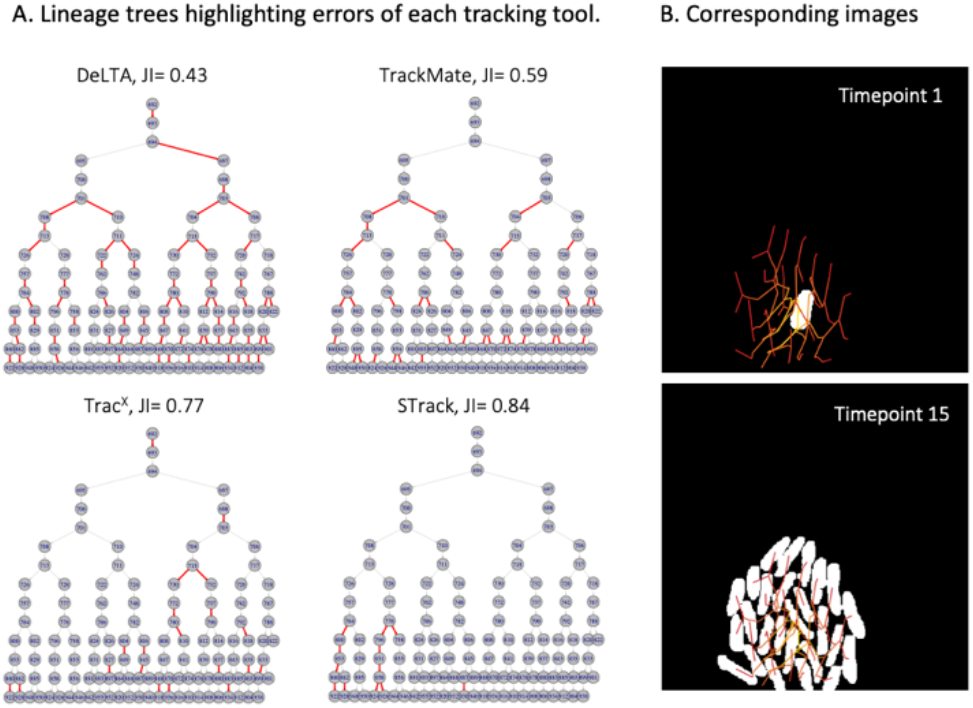
Example of P. putida cell tracking on time-lapse images. A) Lineage trees obtained with the four tracking tools, colored by matches (grey) and mismatches (red) in comparison to manual expert tracking. The calculated Jaccard index (JI) for each comparison is indicated above the corresponding lineage tree. B) Corresponding cell images of the first and last image of the time-lapse, with manual expert-annotated cell masks represented as white cells. Cell tracks are shown in a red line network overlay on the images.

**Fig. 8:**
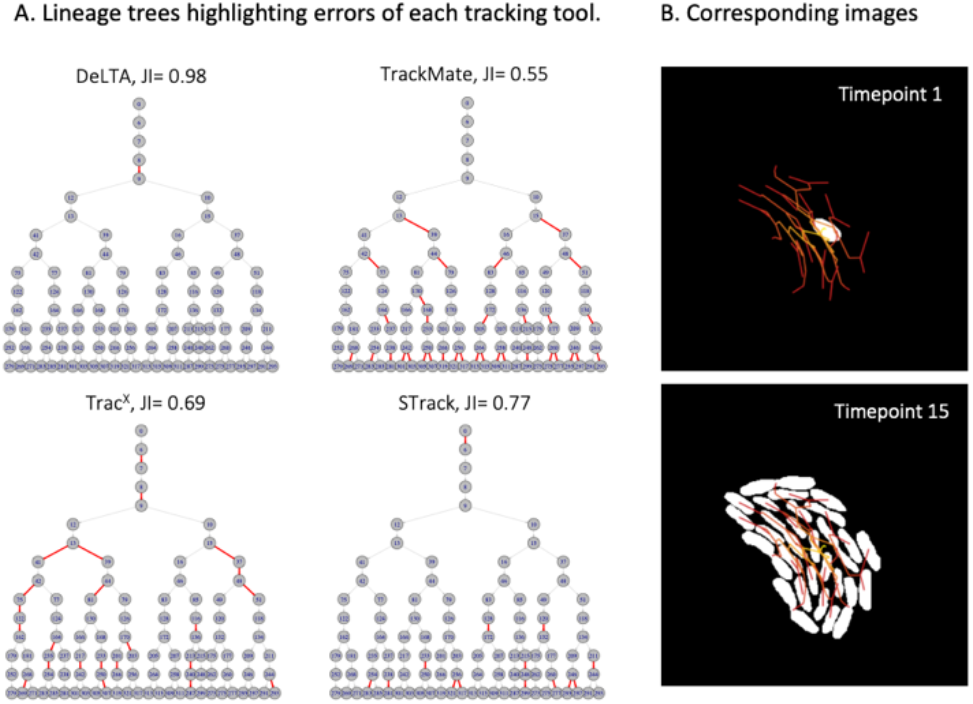
Visual comparison of DeLTA 2.0, TrackMate, Trac^X^ and STrack on a second area of the P.putida time-lapse. A) Compared to Figure 7, the global performance of all tools except DeLTA 2.0 remained relatively similar. B) Corresponding cell images of the first and last time point of the time-lapse, with manual expert-annotated cell masks represented as white cells. Cell tracks are shown in a red line network overlay on the images.

Finally, we assessed the time requirement for tracking as a function of the number of cells per time-lapse. We applied the four tracking tools on datasets containing between 1156 cells and over 6000 cells (Fig. 9, Supplementary table 1). TrackMate was the most efficient tool from a computational point of view, steadily completing the tracking task in a few seconds, regardless of the data size (Fig. 9). The time necessary for Trac^X^ and STrack to complete tracking increased with the number of cells to track, but remained within four minutes for 6000 cells. DeLTA 2.0 was the least efficient of the tools we tested in terms of timing, taking between five minutes in the smallest dataset to 33 minutes in the largest dataset.

**Fig. 9:**
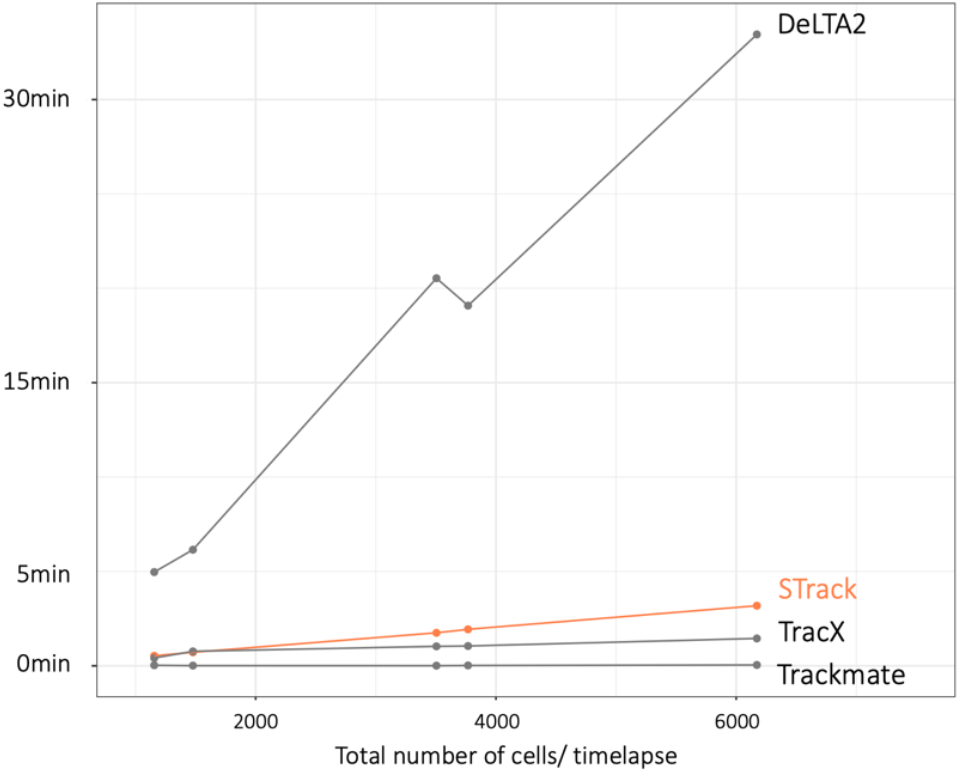
Comparison of the running times of TrackMate, Trac^X^, DeLTA 2.0 and STrack, on five time-lapses containing between 1156 and 6172 cells. All tools but one completed cell tracking in less than one minute on the smallest datasets, and in less than four minutes on the largest dataset. DeLTA 2.0, on the other hand, was significantly slower. It took four more minutes to run on the smallest datasets, and it needed over 30 minutes to track cells in the largest dataset.

Different aspects of STrack, TrackMate, Trac^X^ and DeLTA 2.0 that we consider valuable from a user’s point of view are summarized in Table 1. These concern, notably, the tool’s availability in terms of open access, user-friendliness in terms of the number of parameters to be set and necessity to rename files following a specific syntax, computation speed and accuracy of results. All tools compared in this paper are free, except for Trac^X^ that requires a Matlab license. Trac^X^ also requires specific file naming (as does DeLTA 2.0) and has 50 parameters that can be tuned, which makes it less user-friendly compared to STrack or DeLTA 2.0. TrackMate was the fastest tool we tested, and the highest overall accuracy in the tracking results was obtained with STrack.

**Table 1:** Tracking tool global comparison

| Tool | Open access | Need file renaming | Number of parameters | Computational speed | Accuracy (JI) |
| --- | --- | --- | --- | --- | --- |
| STrack | Yes | No | 2 | Moderate | 0.838 |
| TrackMate | Yes | No | 11 | Fast | 0.696 |
| TracX | No | Yes | 50 | Moderate | 0.803 |
| DeLTa 2.0 | Yes | Yes | 0 | Slow | 0.699 |

## 4 Discussion

Cell tracking in time-lapse images is important because it allows to gain insight in the real-time behaviour of bacteria, on how cells divide, move, or express specific other characteristics (e.g., activation of mobile elements (Sulser et al., [2022]), induction of tailocyns (Vacheron et al. [2021])). Identifying segmented cells and arranging them into lineages by hand or semi manually quickly becomes untractable. Furthermore, high cell numbers are needed for appropriate statistical comparison of cellular behaviours. There is thus a clear need for automated tools that can segment cells correctly and rapidly group those in relevant tracks, with high accuracy. In the tasks we posed ourselves here, namely: deriving cell lineages from dividing cells on surfaces into microcolonies, we found that our tool STrack outperformed three other recently published tracking tools. This was tested on time-lapse image series with six different bacterial strains, five of which had rod-shaped morphologies and the sixth displaying coccoid cells in elongated chains. STrack consistently returned highly accurate results even in suboptimal images, where microcolonies grew close to image borders and eventually out of frame.

STrack was meant to track segmented objects in time-lapses. The quality of segmentation thus plays a crucial role, as mis-segmentations such as missed cells or merged cells will inevitably have a deleterious impact on the tool that tries to link these cells between frames. To avoid such segmentation errors, and to generate inputs that would least influence tracking results for optimal comparison of tracking tools, we decided to segment cell contours manually in all the datasets presented in this paper. The tedious task of segmenting cells manually can however be replaced by powerful automated segmentation tools (Panigrahi et al. [2021], Cutler et al. [2022]).

Many different tools can be used to perform cell tracking, and it can be difficult for a user to decide on tool preference. In this context, STrack can offer a reliable first solution, as it is fast and simple to use, and consistently returns accurate tracking results. For our comparison, we chose not to include interactive tools such as FAST, or Ilastik’s “Tracking with Learning” module (Haubold et al. [2016], Berg et al. [2019]), which help to achieve excellent tracking results but only after thorough training and curation with high demand of user interference. The main reason for their exclusion was the long manual handling to achieve tracking. On the other hand, TrackMate, Trac^X^ and DeLTA 2.0 can be applied on segmented masks directly, and only require several parameters to be set (which is not even necessary in the case of DeLTA 2.0). To have a fair comparison, we applied all four methods (STrack, TrackMate, Trac^X^ and DeLTA 2.0) on the same datasets, with expert-annotated tracks as reference for the evaluation. Across all datasets, we found that STrack’s results were consistently closest to the manual expert annotation, but we acknowledge that some of the other tracking tools were more precise in certain cases, for example in the case of *Rahnella* microcolony growth. It might thus be good practice to test different tracking tools, as their performance might be dataset dependent. As an example, we noticed that the effectiveness of DELTA 2.0 was correlated with the distance of cells to image borders. Making sure to frame cells such that they grow at the centre of time-lapse images, or re-training DeLTA 2.0 specifically with microcolonies that grow on image borders might be solutions to improve its accuracy.

So far, authors of tracking tools have mainly focused on tracking nonmotile or low-motile bacteria, assuming that their close cell-neighbour environment is not changing drastically from one frame to the next. This is also the case of STrack, as one of its limitations is that it will always link bacteria that are in close vicinity, thus making it inappropriate when bacteria start to move around. In order to tackle the much more complex challenge of tracking motile bacteria in time-lapse images, such as bacteria with a predatory behaviour (i.e., *Myxococcus xanthus* or *Lysobacter*, Seef et al. [2021]), a new generation of tracking tools will be required. These tools will need to evaluate all possible cell-to-cell combinations to select the most plausible tracking scenarios at the image scale, as opposed to currently looking in the close vicinity of each cell. Alternatively, or additionally, a new generation of tracking tools will also have to consider each bacteria’s movement vector, to predict plausible bacterial trajectories. This idea has recently been developed at the image scale (Dal Co et al. [2020]), but movement vectors will need to be extracted at the single cell level if we want to reach correct tracking even when bacteria change direction during the time-lapses. This will make the task of cell tracking highly computationally challenging.

Analysing images in an automated way has the advantage to improve reproducibility as compared to manual data extraction. Unfortunately, even the same piece of code can still lead to variable results on different computers due to varying package versions and operating systems. STrack overcomes this issue as it is wrapped in a docker structure, in which the versions of all necessary packages are hard-coded. STrack will thus more easily operate across different systems, facilitating the generation of consistent results, regardless of the date or the computer on which it is launched. With reliably accurate and stable tools such as STrack, we aim to contribute to better detection and quantification of interesting biological cell behaviours from imaging, thus facilitating our acquisition of real-time knowledge on bacterial interactions.

## 5 Data availability

The data underlying this article will be made available on Zenodo.

## 6 Acknowledgements

This work was supported by the Swiss National Science Foundation Sinergia program [grant number CRSII5_189919/1] and by the National Center of Competence in Research NCCR *Microbiomes*.

We kindly thank Andreas Cuny for providing the MATLAB script for Trac^X^ and helping to troubleshoot with the package. We would also like to acknowledge the work of Elvire Sarton-Lohéac, Isaline Guex, Senka Causevic and Maxime Batsch, who helped with manual expert cell segmentation and cell tracking in countless images.

## Algorithm 1 STrack

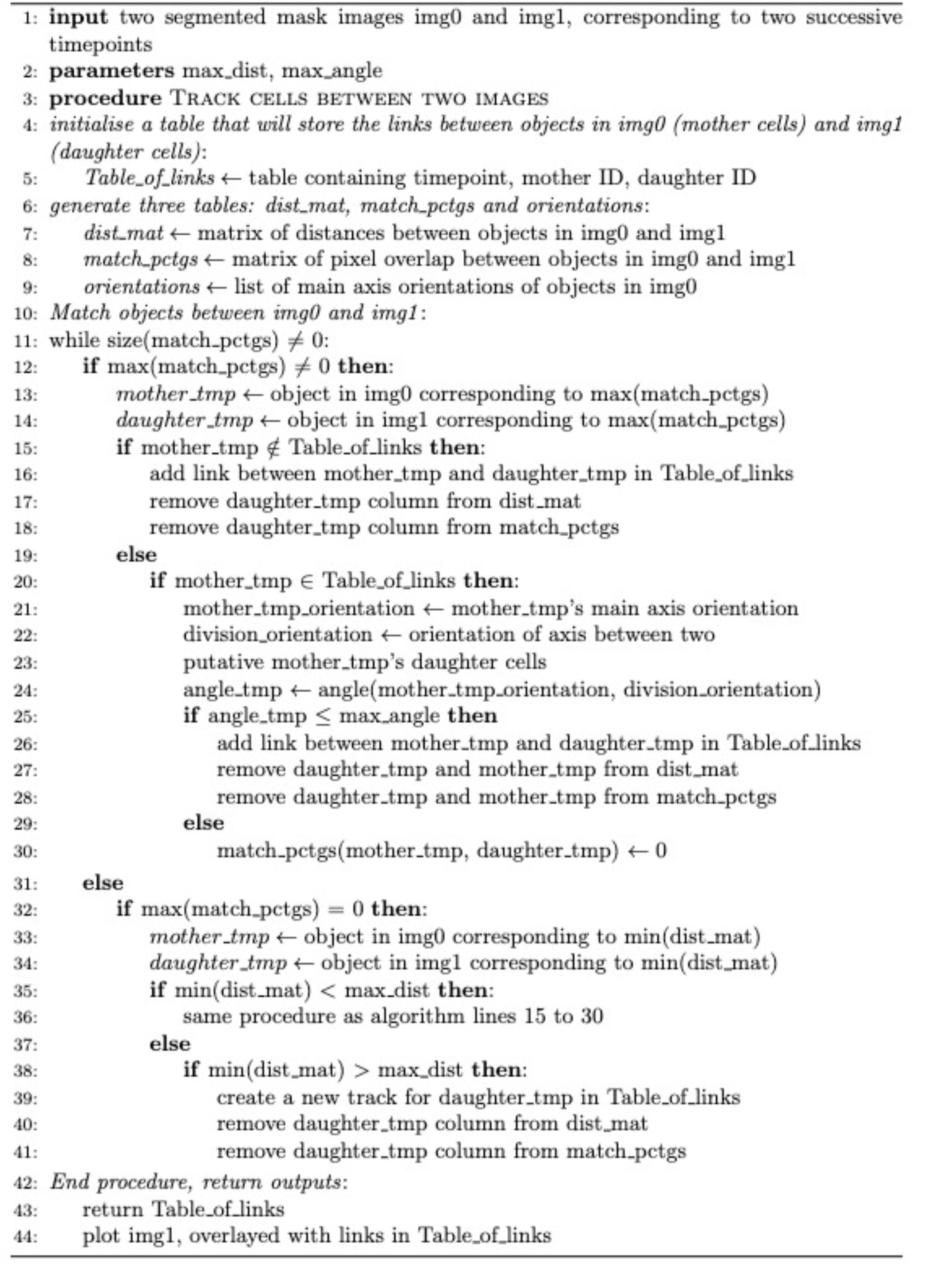

